# Historical specimens and the limits of subspecies phylogenomics in the New World quails (Odontophoridae)

**DOI:** 10.1101/2021.11.18.468700

**Authors:** Jessie F. Salter, Peter A. Hosner, Whitney L. E. Tsai, John E. McCormack, Edward L. Braun, Rebecca T. Kimball, Robb T. Brumfield, Brant C. Faircloth

## Abstract

As phylogenomics focuses on comprehensive taxon sampling at the species and population/subspecies levels, incorporating genomic data from historical specimens has become increasingly common. While historical samples can fill critical gaps in our understanding of the evolutionary history of diverse groups, they also introduce additional sources of phylogenomic uncertainty, making it difficult to discern novel evolutionary relationships from artifacts caused by sample quality issues. These problems highlight the need for improved strategies to disentangle artifactual patterns from true biological signal as historical specimens become more prevalent in phylogenomic datasets. Here, we tested the limits of historical specimen-driven phylogenomics to resolve subspecies-level relationships within a highly polytypic family, the New World quails (Odontophoridae), using thousands of ultraconserved elements (UCEs). We found that relationships at and above the species-level were well-resolved and highly supported across all analyses, with the exception of discordant relationships within the two most polytypic genera which included many historical specimens. We examined the causes of discordance and found that inferring phylogenies from subsets of taxa resolved the disagreements, suggesting that analyzing subclades can help remove artifactual causes of discordance in datasets that include historical samples. At the subspecies-level, we found well-resolved geographic structure within the two most polytypic genera, including the most polytypic species in this family, Northern Bobwhites (*Colinus virginianus*), demonstrating that variable sites within UCEs are capable of resolving phylogenetic structure below the species level. Our results highlight the importance of complete taxonomic sampling for resolving relationships among polytypic species, often through the inclusion of historical specimens, and we propose an integrative strategy for understanding and addressing the uncertainty that historical samples sometimes introduce to phylogenetic analyses.

## Introduction

Phylogenomic studies during the previous two decades have used increasing numbers of loci to resolve relationships at finer and finer taxonomic scales from families (Hackett et al., 2008) to genera (Burleigh et al., 2015) to species (Harvey et al., 2020). Although some of these deeper relationships are still debated (Reddy et al., 2017), the attention of phylogenetics has begun to turn towards resolving relationships at and below the species level (Harvey et al., 2016). Dense sampling at the species and subspecies levels has historically been limited by the cost of generating sequence data and the availability of tissues suitable for DNA extraction. However, improvements in sequencing and laboratory techniques have provided solutions to both problems by reducing the unit cost of sequencing and enabling the collection of genome-scale data from contemporary and historical sources, such as museum specimens (Bi et al., 2013; Derkarabetian et al., 2019; Faircloth et al., 2015; McCormack et al., 2017; Ruane and Austin, 2017; Tsai et al., 2019b). Historical specimens are often used to fill sampling gaps left by rare, endangered, or extinct taxa that lack available tissues, and their inclusion in phylogenomic analyses can dramatically reshape our understanding of the evolutionary history, systematics, and taxonomy of organismal groups (Salter et al., 2020). Yet, for all the opportunities museum specimens offer, they also introduce novel methodological challenges and potential sources of error in downstream phylogenomic analyses, particularly when a study focuses on resolving fine-scale differences among species and subspecies.

For example, previous studies incorporating historical specimens have noted several recurrent issues associated with sample quality that manifest as failures to detect some loci, shorter contigs assembled for detected loci, and DNA damage within assembled contigs (Hosner et al., 2016; Moyle et al., 2016; Oliveros et al., 2019; Salter et al., 2020; Smith et al., 2020; Swanson et al., 2019). These effects can lead to analytical issues like abnormally long branch lengths (McCormack et al., 2012), alternative placements of taxa between concatenated and coalescent analyses (Moyle et al., 2016; Oliveros et al., 2019; Salter et al., 2020), and consistent placement of historical samples as sister to all remaining taxa within certain clades (Moyle et al., 2016; Oliveros, 2015). Discordant topologies that include historical samples are especially vexing because it can be unclear whether legitimate differences arise from more complete taxonomic sampling or whether the incorporation of sequencing and assembly errors from lower quality samples is driving spurious results. Unresolved differences in placement can also leave lingering uncertainty surrounding the evolutionary history of lineages that might be important targets for conservation or additional study (Salter et al., 2020). Although these issues have been noted repeatedly, few studies (Moyle et al., 2016; Oliveros, 2015; Smith et al., 2020) have explored mechanisms for addressing these apparent analytical artifacts.

Here, we use historical and contemporary specimens to reconstruct a subspecies phylogeny of a highly polytypic group of birds, the New World quails (Odontophoridae). New World quails are small (140-170g) terrestrial birds found in forest and grassland habitats from southern Canada to southeastern Brazil and northern Argentina (Brennan et al., 2020; Carroll, 1994). Originally named for the serrated edge of their mandible (from the Greek *odonto*, tooth, *phor,* bearer, i.e. tooth- bearer) (Johnsgard, 1988), New World quails are distinguished by their complex plumage patterns and occasional head ornamentation, ranging from crests to teardrop-shaped plumes to single- feather “spikes.” The family reaches peak diversity in southern Mexico and Central America, where 17 species are found and up to eight species may co-occur (Johnsgard, 1988).

The taxonomic status of the New World quails has long been the subject of debate. Although New and Old World quails have been recognized as distinct clades since the first comprehensive taxonomy of quails and partridges (Ogilvie-Grant, 1893), phenotypic similarities between New and Old World quail species resulted in the description of New World quails as either a tribe (Odontophorini; Verheyen, 1956) or a subfamily (Odontophorinae; Ogilvie-Grant, 1896) within the pheasants (Phasianidae). Based on comparative osteological evidence, Holman (1961) argued New World quails warranted recognition as a distinct family, an idea validated by DNA-DNA hybridization analyses (Sibley and Ahlquist, 1986, 1985) that showed New World quails were more divergent from Old World galliformes than other New World taxa such as turkeys and grouse. More recent molecular studies of Galliformes have confirmed the placement of New World quails as sister to pheasants (Phasianidae; Cox et al., 2007; Hosner et al., 2016; Kimball and Braun, 2014; Wang et al., 2013). Surprisingly, these results also revealed a sister relationship between the clade of New World quails and African (Old World) partridges in the genus *Ptilopachus* (Cohen et al., 2012; Crowe et al., 2006; Hosner et al., 2015), calling into question whether the sister lineage of pheasants consists of only “New World” species. Dating analyses and inferred rates of sequence evolution suggest *Ptilopachus* and the New World quails diverged from an Old World common ancestor 32 Ma, coincident with the existence of the Beringian land bridge between the Nearctic and Palearctic (Hosner et al., 2015). Because referring to Odontophoridae as “New World” quail is inconsistent with the inclusion of *Ptilopachus*, we will refer to the group as “odontophorids”.

Similar to higher level galliform taxonomy, early systematics within odontophorids used comparative osteology (Holman, 1961) and species ecology (Johnsgard, 1973) to describe the relationships among genera. Both classification schemes identified two major clades (Gutiérrez et al., 1983): the *Odontophorus* group, comprising the genera *Odontophorus*, *Rhynchortyx*, *Dactylortyx*, and *Cyrtonyx*; and the *Dendrortyx* group, comprising the genera *Dendrortyx*, *Philortyx*, *Oreortyx*, *Colinus*, and *Callipepla* (Fig. 1A and 1B). Although molecular studies of odontophorids have validated the general membership of each clade, most recent studies suggest the monotypic genus *Rhynchortyx* is sister to both clades (Fig. 1C) while the arrangement of genera within each clade has differed (Cohen et al., 2012; Crowe et al., 2006; Hosner et al., 2015). The most complete molecular phylogeny of odontophorids (Hosner et al., 2015) included sequence data from three mitochondrial and eight nuclear loci from 23 species and recovered strong support across analyses for intergeneric relationships (Fig. 1C).

**Fig. 1.**
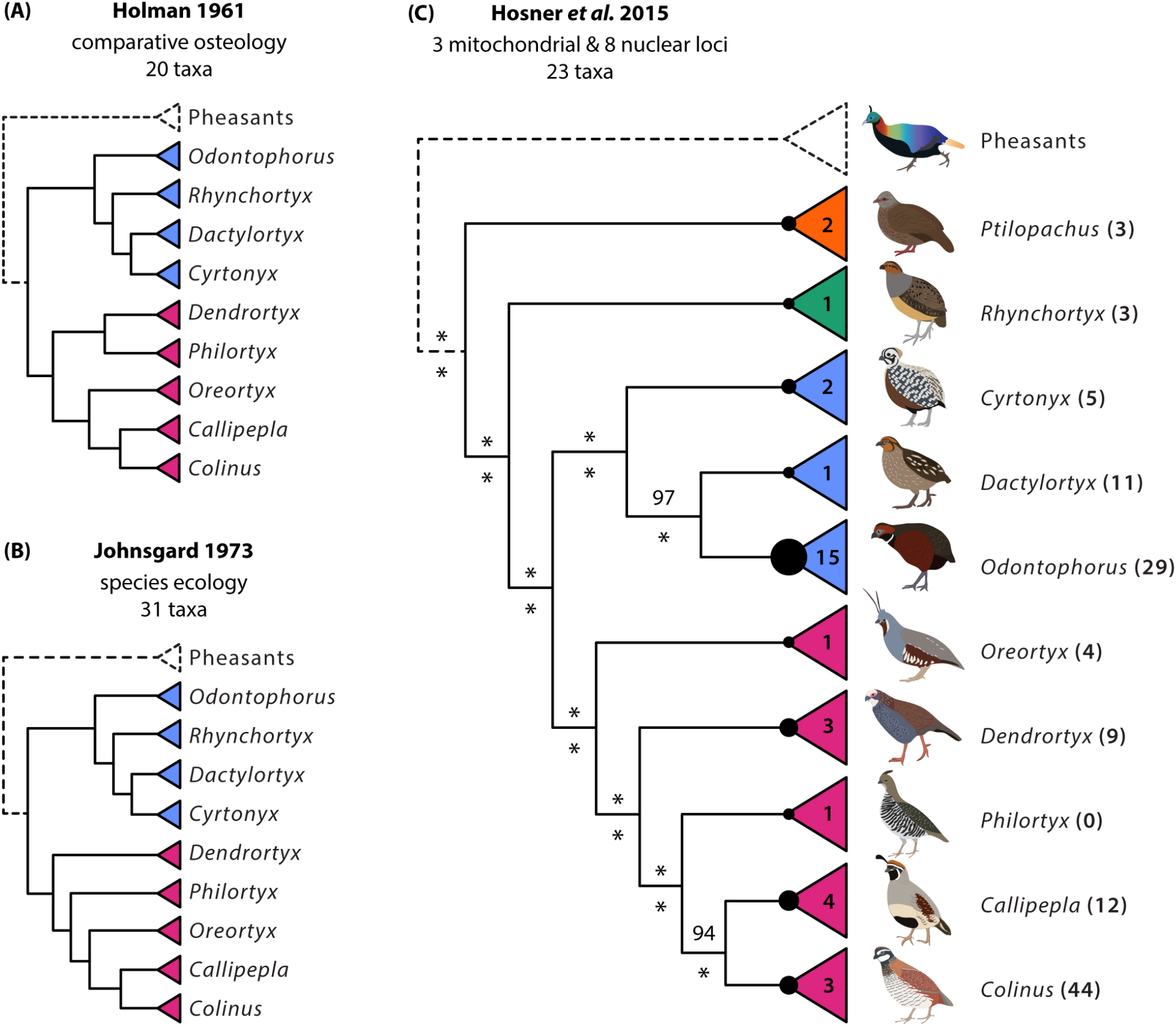
Previous hypotheses of odontophorid relationships based on morphology, ecology, and molecular markers. *Odontophorus* group is shown in blue; *Dendrortyx* group in pink. **(A)** Holman’s (1961) phyletic hypothesis based on comparative osteology. **(B)** Johnsgard’s (1977) phyletic hypothesis based on species ecology. **(C)** Hosner’s (2015) phylogenetic hypothesis based on molecular sequence data. Circles at the base of each genus are scaled to the number of species (shown in triangles); number of subspecies shown in parentheses after each genus. Maximum likelihood (ML) bootstrap support values shown above each node; Bayesian posterior probability shown below. Asterisks indicate 100% ML bootstrapping support / posterior probability.

At the species-level, relationships within odontophorids are less clear. For example, the numbers of odontophorid species and subspecies have fluctuated dramatically through time (Fig. 2), largely due to the difficulty of ascribing consistent taxonomic boundaries to a group that displays remarkable phenotypic variability (Johnsgard, 1988). As a result, different taxonomies recognize anywhere from 27 to 35 species distributed among ten genera (Carroll, 2019; Clements et al., 2019; Dickinson and Remsen, 2013; Johnsgard, 1988). This uncertainty is magnified at the subspecies level, where 126 to 145 subspecies of odontophorids are recognized, primarily based on variation in plumage and disjunctions in geographic ranges (Carroll, 2019; Clements et al., 2019; Dickinson and Remsen, 2013; Johnsgard, 1988). To put this incredible phenotypic diversity in context, odontophorids are more polytypic than 89% of all other bird families (Dickinson and Remsen, 2013), when controlling for family size, including the famously polytypic pheasants. Interestingly, this diversity is not distributed evenly across the family: 13 species of odontophorids are monotypic, while the three species of bobwhites (genus *Colinus*) include 44 subspecies – approximately one-third the total diversity of the entire family (Dickinson and Remsen, 2013). Previous genetic studies with subspecies-level sampling of odontophorids have included only three genera comprising less than half of all subspecies (*Callipepla*, Zink and Blackwell, 1998; *Colinus*, Williford et al., 2016, 2014; and *Dendrortyx*, Tsai et al., 2019a) and, with the exception of *Dendrortyx* (Tsai et al., 2019a), all have used a small number of mitochondrial loci (Williford et al., 2016, 2014; Zink and Blackwell, 1998). Furthermore, within the two most polytypic genera, *Odontophorus* and *Colinus*, different analyses have produced equivocal results, often with low support (Hosner et al., 2015; Williford et al., 2016, 2014). As a result, it is unclear whether the lack of resolution within these relatively young clades (4-5 Ma; Hosner et al., 2015; Williford et al., 2016) reflects real biological signal arising from differences in locus histories due to incomplete lineage sorting or introgression, or whether the lack of resolution is simply due to low power of the small number of loci sampled.

**Fig. 2.**
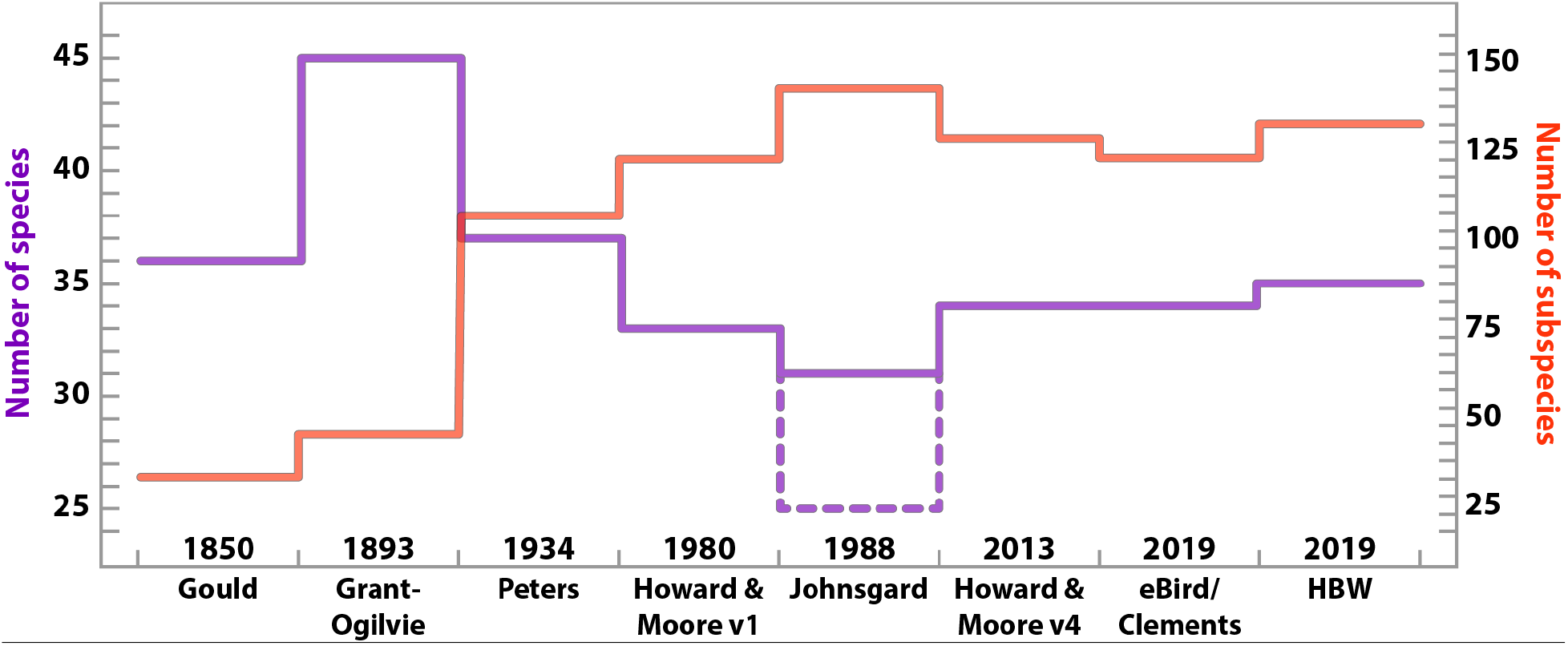
Changes in odontophorid taxonomy through time with different classification schemes (x-axis). The number of species (purple) is shown on the left axis; the number of subspecies (red) is shown on the right axis. Publication year is listed above the x-axis. Dotted line at Johnsgard (1988) shows the number of species when closely-related allopatric species are collapsed to subspecies. HBW = Handbook of the Birds of the World Alive.

Incomplete sampling at the species and subspecies level combined with analyses including few independent loci have limited our understanding at all taxonomic levels within odontophorids and obscured how evolutionary processes may have shaped the remarkable phenotypic diversity observed in this group, but sampling efforts have been limited by lack of access to fresh tissues for many range-restricted and increasingly rare taxa. Sixty-nine percent of odontophorid species have experienced population declines during the past century, and 31% of species are listed as near-threatened or vulnerable by the IUCN (IUCN, 2020). However, historical collections of odontophorids are extensive (>40,000 specimens listed on VertNet.org as of February 2021) due to their popularity as game birds. The extensive availability of historical specimens makes odontophorids an ideal taxonomic group to address some of the larger questions about the role of sample quality in phylogenomic analyses at the species and subspecies level, which we investigate by performing an analysis of ultraconserved elements (UCEs) collected from 42 modern tissues, 83 historical specimens, and six published genomes representing 115 odontophorid taxa (88% of all subspecies) from 83 states/provinces in 22 countries.

## Methods

### Taxonomy

For the sake of clarity throughout the manuscript, we followed version 4 of the Howard and Moore taxonomy (Dickinson and Remsen, 2013), which recognizes 10 genera, 33 species, and 131 subspecies of odontophorids.

### Sampling and DNA extraction

We collected new sequence data from 120 samples, including 78 toepads from historical specimens and 42 tissues (Table 1). To avoid re-sampling historical specimens, we also incorporated published sequence data from five individuals (Tsai et al., 2019b), and we harvested UCE loci from whole genome assemblies (Table 1) for six additional individuals using Phyluce (Faircloth, 2016) following the Phyluce Tutorial III guidelines (Faircloth, 2015). Whenever possible, we sampled two individuals of each monotypic odontophorid species. Our final sampling design included sequence data collected from 42 tissues, 83 toepads (collected between 1906- 1996), and six published genomes spanning 125 ingroup samples corresponding to 115 of the 131 subspecies of odontophorids (88%) and six outgroup species from other families in Galliformes and the sister order Anseriformes (Table 1).

**Table 1.** Sample information and sequencing statistics. Type refers to type of material: T = tissue; S = sequence; for toepads, the year of collection is given.

| Species / subspecies | Museum / Source | Catalog No. | Type | Read Pairs | Contigs | UCEs | Avg. Locus Length | Collection Locality |
| --- | --- | --- | --- | --- | --- | --- | --- | --- |
| *Anas platyrhynchos* | GenBank | SAMN 10245527 | S | - | 4,215 | 4,195 | 695 | - |
| *Anseranas semipalmata* | GenBank | SAMN 12253809 | S | - | 4,099 | 4,081 | 663 | - |
| *Callipepla californica achrustera* | UWBM | 81488 | T | 791,079 | 4,440 | 4,416 | 953 | Baja California Sur, Mexico |
| *Callipepla californica brunnescens* | LSU | B-29626 | T | 1,451,824 | 4,342 | 4,318 | 1,003 | Nevada, USA |
| *Callipepla californica californica* | LSU | B-17959 | T | 1,246,781 | 4,395 | 4,369 | 1,007 | California, USA |
| *Callipepla californica canfieldae* | LSU | B-34531 | T | 488,930 | 4,204 | 4,176 | 968 | California, USA |
| *Callipepla californica catalinensis* | LACM | 19812 | 1941 | 1,901,089 | 4,061 | 4,039 | 322 | Santa Catalina Island, California, USA |
| *Callipepla douglasii bensoni* | LSU | B-59474 | T | 1,004,915 | 4,411 | 4,387 | 1,012 | Sonora, Mexico |
| *Callipepla douglasii douglasii* | UWBM | 81316 | T | 429,620 | 4,364 | 4,340 | 697 | Sinaloa, Mexico |
| *Callipepla gambelii fulvipectus* | UWBM | 90862 | T | 1,156,937 | 4,330 | 4,304 | 1,008 | Sinaloa, Mexico |
| *Callipepla gambelii gambelii* | LSU | B-62399 | T | 1,323,662 | 4,424 | 4,397 | 1,024 | Texas, USA |
| *Callipepla squamata castanogastris* | LSU | B-64166 | T | 1,205,069 | 4,432 | 4,408 | 1,057 | Texas, USA |
| *Callipepla squamata pallida* | UWBM | 77730 | T | 249,076 | 4,011 | 3,993 | 780 | Arizona, USA |
| *Callipepla squamata squamata* | KU | 29970 | 1951 | 879,187 | 3,899 | 3,882 | 311 | San Luis Potosi, Mexico |
| *Chauna torquata* | GenBank | SAMN<br>12253900 | S | - | 4,225 | 4,204 | 711 | - |
| *Colinus cristatus badius* | FMNH | 419219 | 1956 | 5,266,442 | 4,229 | 4,209 | 394 | Cauca, Colombia |
| *Colinus cristatus barnesi* | AMNH | 325203 | 1939 | 1,776,636 | 4,378 | 4,356 | 361 | Barinas, Venezuela |
| *Colinus cristatus bogotensis* | FMNH | 103888 | 1950 | 2,650,689 | 4,307 | 4,290 | 378 | Cundinamarca, Colombia |
| *Colinus cristatus cristatus* | FMNH | 414969 | 1941 | 2,931,306 | 4,292 | 4,275 | 336 | La Guajira, Colombia |
| *Colinus cristatus decoratus* | WFVZ | 22004 | 1968 | 2,270,955 | 4,258 | 4,234 | 430 | Cesar, Colombia |
| *Colinus cristatus dickeyi* | LACM | 80996 | 1964 | 2,189,662 | 3,970 | 3,950 | 333 | Guanacaste, Costa Rica |
| *Colinus cristatus horvathi* | FMNH | 400048 | 1920 | 1,783,542 | 4,175 | 4,158 | 291 | Merida, Venezuela |
| *Colinus cristatus hypoleucus* | LSU | B-52857 | T | 7,580,091 | 4,285 | 4,265 | 482 | Zacapa, Guatemala |
| *Colinus cristatus incanus* | AMNH | 813158 | 1967 | 2,891,436 | 4,161 | 4,145 | 336 | Guatemala |
| *Colinus cristatus leucopogon* | UWBM | 103378 | T | 1,682,655 | 4,379 | 4,353 | 1,105 | Copan, Honduras |
| *Colinus cristatus leucotis* | FMNH | 419663 | 1958 | 3,327,168 | 4,286 | 4,269 | 401 | Huila, Colombia |
| *Colinus cristatus leylandi* | FLMNH | 30517 | 1946 | 1,801,942 | 3,692 | 3,679 | 251 | Francisco Morazan, Honduras |
| *Colinus cristatus littoralis* | USNM | 386764 | 1945 | 4,543,969 | 4,220 | 4,192 | 499 | Magdalena, Colombia |
| *Colinus cristatus mariae* | FMNH | 400209 | 1913 | 1,548,414 | 4,166 | 4,148 | 300 | Chiriqui, Panama |
| *Colinus cristatus mocquerysi* | LACM | 35924 | 1958 | 3,442,312 | 4,041 | 4,023 | 338 | Monagas, Venezuela |
| *Colinus cristatus panamensis* | AMNH | 233102 | 1925 | 2,178,271 | 3,769 | 3,713 | 320 | Veraguas, Panama |
| *Colinus cristatus parvicristatus* | FMNH | 425597 | 1976 | 3,467,525 | 4,327 | 4,302 | 534 | Meta, Colombia |
| *Colinus cristatus sclateri* | FMNH | 371183 | 1970 | 1,405,122 | 4,197 | 4,173 | 329 | Managua, Nicaragua |
| *Colinus cristatus sonnini* | FMNH | 391236 | T | 23,331,061 | 4,494 | 4,467 | 901 | Amapa, Brazil |
| *Colinus nigrogularis caboti* | FLMNH | 18476 | 1973 | 3,517,697 | 4,252 | 4,227 | 430 | Yucatan, Mexico |
| *Colinus nigrogularis nigrogularis* | WFVZ | 21000 | 1969 | 4,618,473 | 4,323 | 4,299 | 528 | Cayo, Belize |
| *Colinus nigrogularis persiccus* | KU | 577 | T | 1,350,192 | 4,395 | 4,368 | 1,054 | Yucatan, Mexico |
| *Colinus virginianus aridus* | LACM | 77830 | 1947 | 1,566,793 | 3,451 | 3,396 | 321 | Tamaulipas, Mexico |
| *Colinus virginianus atriceps* | MLZ | 50749 | 1950 | 3,416,655 | 4,284 | 4,265 | 349 | Guerrero, Mexico |
| *Colinus virginianus coyoleos* | FLMNH | 30512 | 1953 | 2,470,229 | 4,008 | 3,991 | 274 | Oaxaca, Mexico |
| *Colinus virginianus cubanensis* | FMNH | 371181 | 1959 | 2,800,259 | 4,300 | 4,281 | 340 | Pinar del Rio, Cuba |
| *Colinus virginianus floridanus* | LSU | B-48982 | T | 1,321,106 | 4,439 | 4,411 | 1,010 | Florida, USA |
| *Colinus virginianus godmani* | AMNH | 768798 | 1962 | 3,725,131 | 4,263 | 4,238 | 364 | Oaxaca, Mexico |
| *Colinus virginianus graysoni* | USNM | 573690 | 1977 | 13,531,186 | 4,316 | 4,296 | 732 | Nayarit, Mexico |
| *Colinus virginianus harrisoni* | WFVZ | 16994 | 1965 | 2,530,556 | 4,321 | 4,299 | 386 | Oaxaca, Mexico |
| *Colinus virginianus insignis* | FLMNH | 30515 | 1953 | 3,231,160 | 4,150 | 4,135 | 301 | Chiapas, Mexico |
| *Colinus virginianus maculatus* | FMNH | 415417 | 1941 | 2,937,623 | 4,213 | 4,195 | 352 | Tamaulipas, Mexico |
| *Colinus virginianus minor* | WFVZ | 8855 | 1962 | 4,347,086 | 4,298 | 4,281 | 431 | Chiapas, Mexico |
| *Colinus virginianus nigripectus* | WFVZ | 7776 | 1959 | 3,201,973 | 4,313 | 4,292 | 413 | Morelos, Mexico |
| *Colinus virginianus pectoralis* | KU | 23722 | 1946 | 2,449,134 | 4,270 | 4,252 | 312 | Veracruz, Mexico |
| *Colinus virginianus ridgwayi* | YPM | 70184 | 1951 | 3,840,880 | 4,266 | 4,245 | 420 | Sonora, Mexico |
| *Colinus virginianus salvini* | LACM | 80966 | 1971 | 2,304,983 | 4,230 | 4,207 | 389 | Aviary |
| *Colinus virginianus taylori* | LSU | B-62466 | T | 1,375,447 | 4,352 | 4,329 | 1,011 | Texas, USA |
| *Colinus virginianus texanus* | LSU | B-54856 | T | 596,233 | 4,453 | 4,427 | 948 | Texas, USA |
| *Colinus virginianus thayeri* | KU | 45784 | 1964 | 3,094,952 | 4,262 | 4,242 | 417 | Guerrero, Mexico |
| *Colinus virginianus virginianus* | GenBank | SAMN 09044008 | S | - | - | 4,573 | 1,120 | Louisiana, USA |
| *Coturnix japonica* | FMNH | 463188 | T | 1,080,202 | 4,318 | 4,292 | 884 | Captive |
| *Cyrtonyx montezumae mearnsi* | FLMNH | 45132 | T | 1,450,779 | 4,380 | 4,358 | 1,009 | Arizona, USA |
| *Cyrtonyx montezumae merriami* | AMNH | 804738 | 1906 | 1,813,431 | 3,558 | 3,545 | 258 | Veracruz, Mexico |
| *Cyrtonyx montezumae montezumae* | MLZ | 22602 | 1939 | 3,447,980 | 4,177 | 4,162 | 338 | Guanajuato, Mexico |
| *Cyrtonyx montezumae rowleyi* | WFVZ | 19138 | 1967 | 4,072,474 | 4,330 | 4,306 | 495 | Oaxaca, Mexico |
| *Cyrtonyx montezumae sallei* | AMNH | 778476 | 1961 | 3,138,779 | 4,032 | 4,008 | 401 | Jalisco, Mexico |
| *Cyrtonyx ocellatus* | FLMNH | 4315 | 1946 | 2,521,812 | 3,934 | 3,919 | 289 | Morazan, Honduras |
| *Cyrtonyx ocellatus* | MLZ | 56792 | 1954 | 2,883,032 | 4,278 | 4,260 | 350 | Chiapas, Mexico |
| *Dactylortyx thoracicus chiapensis* | FLMNH | 30054 | 1947 | 2,970,682 | 3,864 | 3,853 | 315 | Chiapas, Mexico |
| *Dactylortyx thoracicus conoveri* | FMNH | 412668 | 1937 | 3,021,859 | 3,974 | 3,958 | 371 | Olancho, Honduras |
| *Dactylortyx thoracicus devius* | WFVZ | 4577 | 1959 | 3,177,198 | 3,995 | 3,969 | 540 | Jalisco, Mexico |
| *Dactylortyx thoracicus dolichonyx* | MLZ | 36952 | 1943 | 2,611,157 | 4,110 | 4,091 | 367 | Chiapas, Mexico |
| *Dactylortyx thoracicus fuscus* | FMNH | 412669 | 1937 | 3,021,895 | 4,167 | 4,150 | 340 | Tegucigalpa, Honduras |
| *Dactylortyx thoracicus melodus* | MLZ | 45995 | 1947 | 2,388,679 | 4,306 | 4,290 | 405 | Guerrero, Mexico |
| *Dactylortyx thoracicus paynteri* | YPM | 13023 | 1951 | 3,965,704 | 3,964 | 3,941 | 467 | Quintana Roo, Mexico |
| *Dactylortyx thoracicus pettingilli* | MLZ | 38766 | 1943 | 4,509,113 | 4,065 | 4,046 | 363 | San Luis Potosi, Mexico |
| *Dactylortyx thoracicus sharpei* | FMNH | 215965 | 1951 | 2,687,978 | 3,961 | 3,932 | 434 | Yucatan, Mexico |
| *Dendrortyx barbatus* | UNAM | 24556 | T | 974,224 | 4,321 | 4,297 | 952 | Mexico |
| *Dendrortyx leucophrys hypospodius* | WFVZ | 53075 | 1996 | 5,398,898 | 3,370 | 3,345 | 384 | San Jose, Costa Rica |
| *Dendrortyx leucophrys leucophrys* | MLZ | 26402 | 1934 | 557,857 | 3,166 | 3,155 | 373 | Francisco Morazan, Honduras |
| *Dendrortyx macroura diversus* | WFVZ | 4576 | 1959 | 5,195,088 | 3,989 | 3,969 | 536 | Jalisco, Mexico |
| *Dendrortyx macroura griseipectus* | MLZ | 30358 | 1941 | 420,216 | 3,067 | 3,054 | 346 | Mexico, Mexico |
| *Dendrortyx macroura inesperatus* | MLZ | 46105 | 1947 | 500,914 | 3,158 | 3,147 | 404 | Guerrero, Mexico |
| *Dendrortyx macroura macroura* | MLZ | 46880 | 1947 | 767,043 | 3,470 | 3,458 | 424 | Puebla, Mexico |
| *Dendrortyx macroura oaxacae* | MLZ | 65301 | 1965 | 472,917 | 2,932 | 2,916 | 356 | Oaxaca, Mexico |
| *Gallus gallus* | GenBank | SAMN 00000795 | S | - | 4,314 | 4,296 | 698 | - |
| *Odontophorus atrifrons atrifrons* | FMNH | 405057 | 1927 | 3,013,730 | 4,154 | 4,139 | 323 | Magdalena, Colombia |
| *Odontophorus atrifrons navai* | USNM | 372441 | 1942 | 1,692,609 | 4,004 | 3,989 | 277 | La Guajira, Colombia |
| *Odontophorus atrifrons variegatus* | LACM | 40827 | 1962 | 3,998,487 | 4,222 | 4,202 | 395 | Santander, Colombia |
| *Odontophorus balliviani* | KU | 21232 | T | 1,032,001 | 4,469 | 4,442 | 962 | Puno, Peru |
| *Odontophorus balliviani* | YPM | 38340 | 1956 | 2,516,854 | 4,227 | 4,206 | 450 | Cochabamba, Bolivia |
| *Odontophorus capueira capueira* | YPM | 66400 | 1961 | 5,728,379 | 4,238 | 4,216 | 486 | Misiones, Argentina |
| *Odontophorus capueira plumbeicollis* | FMNH | 395742 | T | 6,929,116 | 4,363 | 4,337 | 984 | Brazil |
| *Odontophorus columbianus* | FMNH | 408566 | 1956 | 2,601,073 | 3,486 | 3,475 | 254 | Carabobo, Venezuela |
| *Odontophorus columbianus* | USNM | 483540 | 1955 | 543,405 | 3,299 | 3,242 | 273 | Venezuela |
| *Odontophorus dialeucos* | USNM | 484271 | 1964 | 4,104,166 | 4,312 | 4,293 | 435 | Panama |
| *Odontophorus dialeucos* | USNM | 484272 | 1964 | 4,384,715 | 4,291 | 4,270 | 408 | Panama |
| *Odontophorus erythrops erythrops* | LSU | B-7868 | T | 1,350,803 | 4,435 | 4,412 | 1,040 | El Oro, Ecuador |
| *Odontophorus erythrops parambae* | LSU | B-1412 | T | 1,230,732 | 4,440 | 4,416 | 997 | Darien, Panama |
| *Odontophorus gujanensis buckleyi* | ANSP | 186791 | T | 931,813 | 4,474 | 4,459 | 441 | Sucumbios, Ecuador |
| *Odontophorus gujanensis castigatus* | FMNH | 408884 | 1931 | 2,395,695 | 3,255 | 3,217 | 276 | Puntarenas, Costa Rica |
| *Odontophorus gujanensis gujanensis* | LSU | B-35494 | T | 694,040 | 4,434 | 4,413 | 862 | Mato Grosso, Brazil |
| *Odontophorus gujanensis marmoratus* | LACM | 40594 | 1961 | 4,252,551 | 4,205 | 4,185 | 432 | Magdalena, Colombia |
| *Odontophorus gujanensis medius* | AMNH | DOT-14437 | T | 764,410 | 4,466 | 4,447 | 995 | Amazonas, Brazil |
| *Odontophorus gujanensis pachyrhynchus* | LSU | B-11238 | T | 1,006,320 | 4,437 | 4,417 | 990 | Ucayali, Peru |
| *Odontophorus gujanensis rufogularis* | LSU | B-40408 | T | 576,970 | 4,283 | 4,260 | 820 | Loreto, Peru |
| *Odontophorus gujanensis simonsi* | LSU | B-15196 | T | 707,114 | 3,272 | 3,258 | 813 | Santa Cruz, Bolivia |
| *Odontophorus guttatus* | FLMNH | 4312 | 1949 | 2,223,632 | 3,983 | 3,973 | 271 | Oaxaca, Mexico |
| *Odontophorus hyperythrus* | FMNH | 418804 | 1951 | 2,520,157 | 4,225 | 4,206 | 336 | Antioquia, Colombia |
| *Odontophorus hyperythrus* | YPM | 54500 | 1957 | 3,895,349 | 4,291 | 4,273 | 423 | Cauca, Colombia |
| *Odontophorus leucolaemus* | USNM | 652312 | T | 998,587 | 4,484 | 4,464 | 981 | Chiriqui, Panama |
| *Odontophorus melanonotus* | FMNH | 414494 | 1940 | 2,685,292 | 4,249 | 4,234 | 356 | Pinchincha, Ecuador |
| *Odontophorus melanonotus* | FMNH | 419577 | 1957 | 3,128,562 | 4,184 | 4,170 | 375 | Narino, Colombia |
| *Odontophorus melanotis melanotis* | FMNH | 412694 | 1966 | 1,864,623 | 4,254 | 4,239 | 332 | Bocas del Toro, Panama |
| *Odontophorus melanotis verecundus* | LSU | 31069 | 1963 | 945,462 | 4,340 | 4,314 | 314 | Olancho, Honduras |
| *Odontophorus speciosus loricatus* | FMNH | 433045 | T | 879,664 | 4,448 | 4,418 | 948 | Cuzco, Peru |
| *Odontophorus speciosus soderstromii* | WFVZ | 42447 | 1987 | 3,230,882 | 3,927 | 3,907 | 564 | Morona Santiago, Ecuador |
| *Odontophorus speciosus speciosus* | LSU | B-43621 | T | 902,333 | 3,734 | 3,715 | 822 | San Martin, Peru |
| *Odontophorus stellatus* | LSU | B-9314 | T | 709,643 | 3,149 | 3,133 | 807 | Pando, Bolivia |
| *Odontophorus stellatus* | LSU | B-27503 | T | 989,421 | 3,564 | 3,547 | 801 | Loreto, Peru |
| *Odontophorus strophium* | AMNH | 176521 | 1920 | 2,394,953 | 4,178 | 4,161 | 320 | Bogota, Colombia |
| *Odontophorus strophium* | AMNH | 181791 | 1923 | 1,846,844 | 3,935 | 3,917 | 278 | Cundinamarca, Colombia |
| *Oreortyx pictus confinis* | LSU | B-59654 | T | 1,130,780 | 3,944 | 3,926 | 773 | Baja California, Mexico |
| *Oreortyx pictus pictus* | UWBM | 66924 | T | 757,813 | 4,451 | 4,429 | 950 | Washington, USA |
| *Oreortyx pictus plumifer* | UWBM | 86733 | T | 824,244 | 4,465 | 4,440 | 1,010 | California, USA |
| *Oreortyx pictus russellii* | LSU | B-6421 | T | 732,203 | 3,303 | 3,286 | 837 | California, USA |
| *Oxyura jamaicensis* | GenBank | SAMN<br>04270837 | S | - | 4,274 | 4,250 | 656 | - |
| *Philortyx fasciatus* | UNAM | 27429 | T | 793,648 | 4,495 | 4,470 | 986 | Mexico |
| *Ptilopachus nahani* | LACM | 67857 | 1967 | 3,917,333 | 4,206 | 4,188 | 398 | Western Region, Uganda |
| *Ptilopachus nahani* | LACM | 75763 | 1970 | 5,225,360 | 4,164 | 4,141 | 472 | Western Region, Uganda |
| *Ptilopachus petrosus brehmi* | LACM | 41663 | 1960 | 7,239,213 | 3,982 | 3,960 | 390 | Moyen-Chari, Chad |
| *Ptilopachus petrosus florentiae* | LACM | 80630 | 1965 | 6,430,092 | 4,276 | 4,256 | 438 | Rift Valley, Kenya |
| *Ptilopachus petrosus major* | FMNH | 68965 | 1927 | 5,120,149 | 3,872 | 3,847 | 323 | Gojjam, Ethiopia |
| *Rhynchortyx cinctus australis* | LSU | B-29992 | T | 546,267 | 2,723 | 2,707 | 792 | Esmeraldas, Ecuador |
| *Rhynchortyx cinctus cinctus* | LSU | B-1389 | T | 1,145,554 | 3,907 | 3,890 | 834 | Darien, Panama |

We extracted total DNA from tissues using a Qiagen DNeasy Blood & Tissue Kit following the manufacturer’s instructions, and we extracted total DNA from toepads of historical museum specimens using a phenol-chloroform protocol (Tsai et al., 2019b).

### Sequence capture and next-generation sequencing

We prepared genomic libraries from all DNA extracts and performed target enrichment of ultraconserved elements (UCEs; Faircloth et al., 2012) from genomic libraries following the protocol outlined in Salter et al. (2020). In brief, we sheared tissue samples using a QSonica ultrasonicator to a peak size distribution of 400 to 600 bp. We did not shear toepad samples because they already had a peak size distribution of 100 to 300 bp due to DNA degradation (McCormack et al., 2015). We prepared dual-indexed genomic libraries of each sample using the KAPA Hyper Prep library preparation kit (F. Hoffman-LaRoche AG, Basel, Switzerland) and custom indexes (Glenn et al., 2019). We combined the libraries into fourteen pools containing between six and eight samples for enrichment, and we kept tissues and toepads in separate pools. We enriched each library pool for 5,060 UCE loci using a MYbaits_Tetrapods-UCE-5K kit (Daicel Arbor Biosciences, Ann Arbor, MI) following a protocol modified from Faircloth et al. (2012) (Faircloth et al., 2018). After enrichment, we performed 16 cycles of PCR recovery. To remove adapter-dimers, we processed each enriched pool with a Qiagen GeneRead Size Selection Kit, which removes fragments below 150 bp. We then ran post-enrichment pools on a Bioanalyzer to verify peak size distributions and ensure the absence of adapter-dimers. Finally, we quantified pools free of adapter-dimers using the KAPA qPCR quantification kit, and we combined pools at equimolar ratios prior to collecting sequence data using two lanes of 150-bp paired-end (PE150) sequencing on an Illumina HiSeq 3000 (Oklahoma Medical Research Foundation, Oklahoma City, OK).

### Bioinformatic processing, assembly, and alignment of UCEs

After receiving demultiplexed reads from the sequencing facility, we used illumiprocessor (Faircloth, 2013), a wrapper around Trimmomatic (Bolger et al., 2014), to remove adapter sequences from the data and trim raw reads for quality. We followed this same procedure to incorporate reads from the five toepad samples sequenced by Tsai et al. (2019a). Because some libraries received a larger number of FASTQ reads than others, we used seqtk (Li, 2012) to randomly downsample libraries having more than 1.5 million cleaned read pairs (i.e., 3 million reads, in total). We then assembled the data using itero v1.1.2 (Faircloth, 2018), a guided iterative assembly approach designed to improve assembly of target enrichment data. To start the assembly process, itero uses bwa (Li and Durbin, 2009) to seed reads with a reference sequence and assemble loci using SPAdes (Bankevich et al., 2012); each subsequent round of assembly uses the assembled contigs from the previous iteration as the seed. We performed five iterations of assembly with itero using the UCE probe sequences we targeted during enrichment as the initial seed, and after five rounds of assembly we discarded contigs with less than 5x coverage. To check assembled libraries for the correct species identification and potential contamination, we ran the phyluce program match-contigs-to-barcodes (Faircloth, 2016) using a *Colinus virginianus* COI sequence (NCBI GenBank DQ432859.1) as a reference. We then input extracted contigs that matched the reference COI to NCBI BLAST (Johnson et al., 2008) to compare the extracted sequences to those present in NCBI GenBank, confirm the identity of each sample, and check for any contaminating (different species identity) COI sequences. Following assembly, we used phyluce v1.6.7 (Faircloth, 2016) to export the UCE loci into a database, from which we created an incomplete matrix of loci across all samples. We aligned the incomplete matrix with mafft v7.407 (Katoh and Standley, 2013) using default parameters and internally trimmed the alignment with the -automated1 option in trimAl v1.4.rev15 (Capella-Gutiérrez et al., 2009), before creating a matrix of loci where every locus had at least 75% taxon occupancy.

### UCE phylogenies with concatenated and coalescent methods

After concatenating loci in the 75% complete data matrix, we used raxml-ng v0.9.0 (Kozlov et al., 2019) to estimate a maximum likelihood (ML) tree from the unpartitioned data. We estimated 20 ML trees, selected the tree that best fit the data, and we estimated branch support on the best fitting tree by bootstrapping with the autoMRE function, which checks for convergence every 50 bootstraps. The analysis converged after 100 bootstrap replicates, and we reconciled the best ML tree with the bootstrap replicates using raxml-ng. We collapsed nodes with <70% bootstrap support (Hillis and Bull, 1993).

To account for heterogeneous gene/locus histories in our UCE data, we also used a coalescent-based approach to estimate a species tree. Specifically, we selected SVDquartets (Chifman and Kubatko, 2014) for these analyses because our dataset included many historical samples, which have fewer loci and shorter contigs than tissue samples (Table 1), and SVDquartets is less sensitive to these types of missing data than gene tree reconciliation methods (Hosner et al., 2016; Moyle et al., 2016; Oliveros et al., 2019; Salter et al., 2020; Sayyari et al., 2017). To infer the SVDquartets tree, we used PAUP* v4.0a166 (Swofford, 2002) to evaluate all quartets by singular value decomposition and perform bootstrap analysis (svdq evalq=all bootstrap nthreads=12). We collapsed nodes with <70% bootstrap support (Hillis and Bull, 1993).

### Subset phylogenies with concatenated and coalescent methods

We observed several inconsistencies between the concatenated ML topology and the SVDquartets topology (hereafter “complete” datasets) within the *Odontophorus* and *Colinus* clades that we hypothesized were spurious results caused by the inclusion of low-quality historical samples, which can have a higher noise to signal ratio. Specifically, we suspected that strongly conflicting results were caused by the “toepad effect” where short, low-quality UCE contigs assembled from toepad DNA extracts are sometimes resolved as sister to remaining taxa within a clade. Because sequence variability and phylogenetic informativeness increase with distance from the core conserved UCE region (Faircloth et al., 2012) and many toepad samples lack these variable flanking regions, we suspect that toepad samples in other portions of the tree may be responsible for the toepad effect by pulling problematic taxa towards them due to the degree of sequence similarity shared between the relatively short, core UCE regions that are commonly enriched from toepads. To investigate these effects, we subsampled the concatenated dataset output by trimAl to produce two subclades, which we rooted on the most closely related taxa that were stable across the complete ML and SVDquartets analyses: (1) an *Odontophorus* group, rooted on *O. balliviani* and *O. atrifrons*, and (2) a *Colinus* group, which we rooted on *Callipepla sp.* To subsample the trimAl data matrix, we used GNU Grep with regular expressions, and we inferred concatenated and coalescent-based trees using raxml-ng and SVDquartets with parameters that were identical to those applied to the entire concatenated matrix. In both trees, we collapsed nodes with <70% bootstrap support (Hillis and Bull, 1993).

## Results

### Recovery of UCEs

After demultiplexing and trimming the raw reads, we obtained an average of 1.8 million read pairs (range 249,076 - 23,331,061) for tissue samples and 3 million read pairs for toepad samples (range 420,216 - 13,531,186) (Table 1). After downsampling sequence files, we assembled an average of 4,212 (4,126-4,368 95 CI) contigs for tissue samples and 4,043 (3,971-4,115 95 CI) contigs for toepad samples. From the assembled contigs, we identified an average of 4,087 UCE loci, which was consistent across sample types (Table 1); however, the average contig length of the loci differed between sample types: 912 bp (870-954 bp 95 CI) for tissues; 378 bp (360-396 bp 95 CI) for toepads; and 757 bp (614-900 bp 95 CI) for contigs extracted from genome sequences (Table 2). We enriched a total of 3,884 UCE loci shared by at least 98 ingroup and outgroup taxa, which we concatenated into a 75% complete data matrix that comprised 2,005,421 total characters and included 274,886 parsimony informative sites (13.7%).

**Table 2.** Summary statistics of sequencing output by sample type. Values represent mean ± 95% confidence interval. Toepads were collected between 1906-1996.

| Sample type | No. of samples | Avg. clean read pairs | Avg. contigs | Avg. UCE loci | Avg. contig length (bp) | Avg. collection year |
| --- | --- | --- | --- | --- | --- | --- |
| Tissues | 42 | 1,802,937 ± 1,111,737 | 4,212 ± 127 | 4,190 ± 127 | 912 ± 42 | - |
| Toepads | 83 | 3,099,816 ± 385,186 | 4,043 ± 336 | 4,023 ± 73 | 378 ± 18 | 1954 ± 3 |
| Sequences | 6 | - | 4,225 ± 71 | 4,267 ± 133 | 757 ± 143 | - |

### Concatenated UCE phylogeny

The ML tree we inferred from 3,884 concatenated UCE loci was well resolved and strongly supported for most generic and species-level relationships (Fig. 3A; see Supplementary Fig. S1 for branch lengths). Consistent with previous molecular phylogenies (Cohen et al., 2012; Crowe et al., 2006; Hosner et al., 2015), we resolved *Ptilopachus* as sister to all New World species, and, within the New World clade, we resolved the monotypic genus *Rhynchortyx* as sister to the *Odontophorus* and *Dendrortyx* groups.

**Fig. 3.**
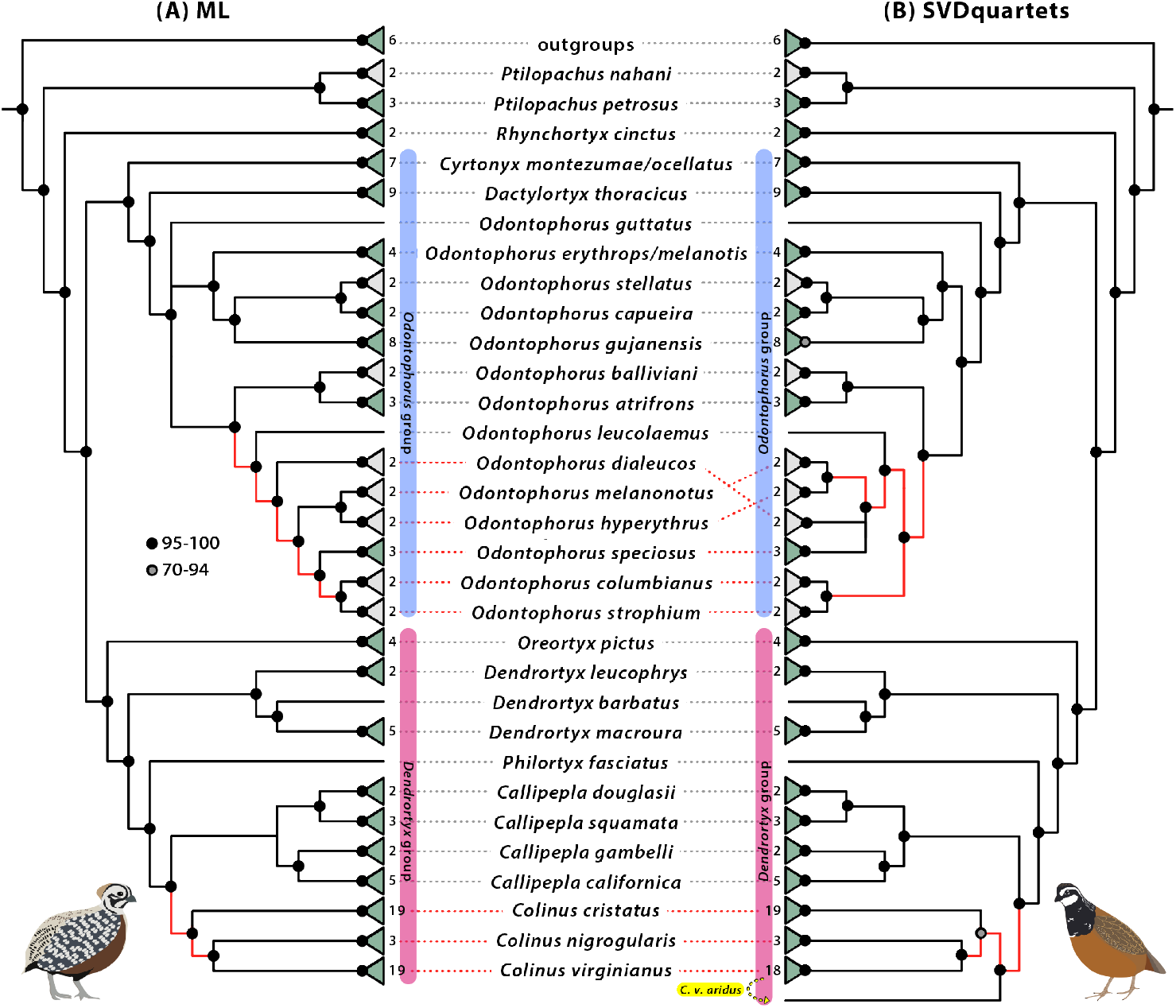
Cladogram of species-level relationships of 125 odontophorid taxa inferred with **(A)** maximum likelihood (ML) analysis and **(B)** SVDquartets analysis of 3,884 nuclear ultraconserved element loci. Gray triangles depict multiple individuals of the same monotypic species; green triangles depict collapsed subspecies relationships. Numbers next to terminal tips indicate the number taxa in collapsed group. Rounded boxes bracketing species names correspond to species groups in Fig. 1. Black circles indicate nodes with >95% bootstrap support; gray circles = 70-94% bootstrap support; branches with <70% bootstrap have been collapsed. Red branches indicate conflicting relationships; yellow highlighting and yellow arrow indicate placement of *C. v. aridus* as sister to remaining *Colinus* in the SVDquartets topology. For a subspecies-level comparison of these trees, see Supplementary Fig. S3. For branch lengths and the uncollapsed ML tree, see Supplementary Fig. S1; for the uncollapsed SVDquartets tree, see Supplementary Fig. S2.

Within the *Odontophorus* group, we resolved the branching order as *Cyrtonyx*, *Dactylortyx*, and *Odontophorus*, and within *Odontophorus*, we resolved the northernmost species, *O. guttatus,* as sister to two clades: one comprising five predominantly lowland tropical forest species (*O. stellatus, O. capueira, O. erythrops, O. melanotis,* and *O. gujanensis*) and one comprising nine montane-associated species (*O. balliviani*, *O. atrifrons*, *O. leucolaemus, O. dialeucos, O. melanonotus*, *O. hyperythrus*, *O. speciosus, O. columbianus*, and *O. strophium*), although support for the branch uniting these two clades was low. We also observed two instances of non-monophyletic species that we detected as a result of subspecies-level sampling: *Cyrtonyx ocellatus* was nested within *C. montezumae*, and *Odontophorus melanotis* was nested within *O. erythrops* (Fig. 3A, Supplementary Fig. S1).

Within the *Dendrortyx* group, we resolved *Oreortyx*, *Dendrortyx*, *Philortyx*, and *Callipepla + Colinus* as successive sister groups, and species-level relationships were consistent with previous studies of each genus (Tsai et al., 2019a; Williford et al., 2016; Zink and Blackwell, 1998).

At the subspecies-level, resolution was more variable across the tree, although results were generally consistent with the broad biogeographic patterns of each species’ distribution (Supplementary Fig. S1). For example, within *Cyrtonyx*, the ML analysis resolved two well-supported clades: one comprising the three subspecies of *C. montezumae* from the southwestern U.S. and northern Mexico (*C. m. mearnsi*, *C. m. merriaimi*, *C. m. montezumae*) and a second clade comprising *C. ocellatus* of southern Mexico and Central America along with the two Oaxacan subspecies of *C. montezumae* (*C. m. sallei*, *C. m. rowleyi*), which are more similar in plumage to *C. ocellatus* and are sometimes considered a separate species named *C. sallei (Carroll, 2019)*. Similarly, we recovered a south-north grade among the two subspecies of *O. erythrops* in Ecuador and Colombia and the two subspecies of *O. melanotis* in Central America, consistent with previous treatments of all four taxa as subspecies of *O. erythrops* (Johnsgard, 1988). Within *Colinus cristatus*, the ML analysis strongly supported two clades: one comprising 13 subspecies of eastern Panama and northern South America and a sister clade comprising the six Central American subspecies of the *C. [cristatus] leucopogon* group (Fig. 4E, Supplementary Fig. S1), which are sometimes treated as a separate species (Carroll, 2019; Johnsgard, 1988). We observed a similar pattern within *C. virginianus*, the most polytypic odontophorid species: a well-supported split between eight subspecies north of Mexico’s Transvolcanic belt and eleven subspecies south of this barrier, although we were generally unable to resolve phylogenetic relationships among subspecies within either of these clades.

**Fig. 4.**
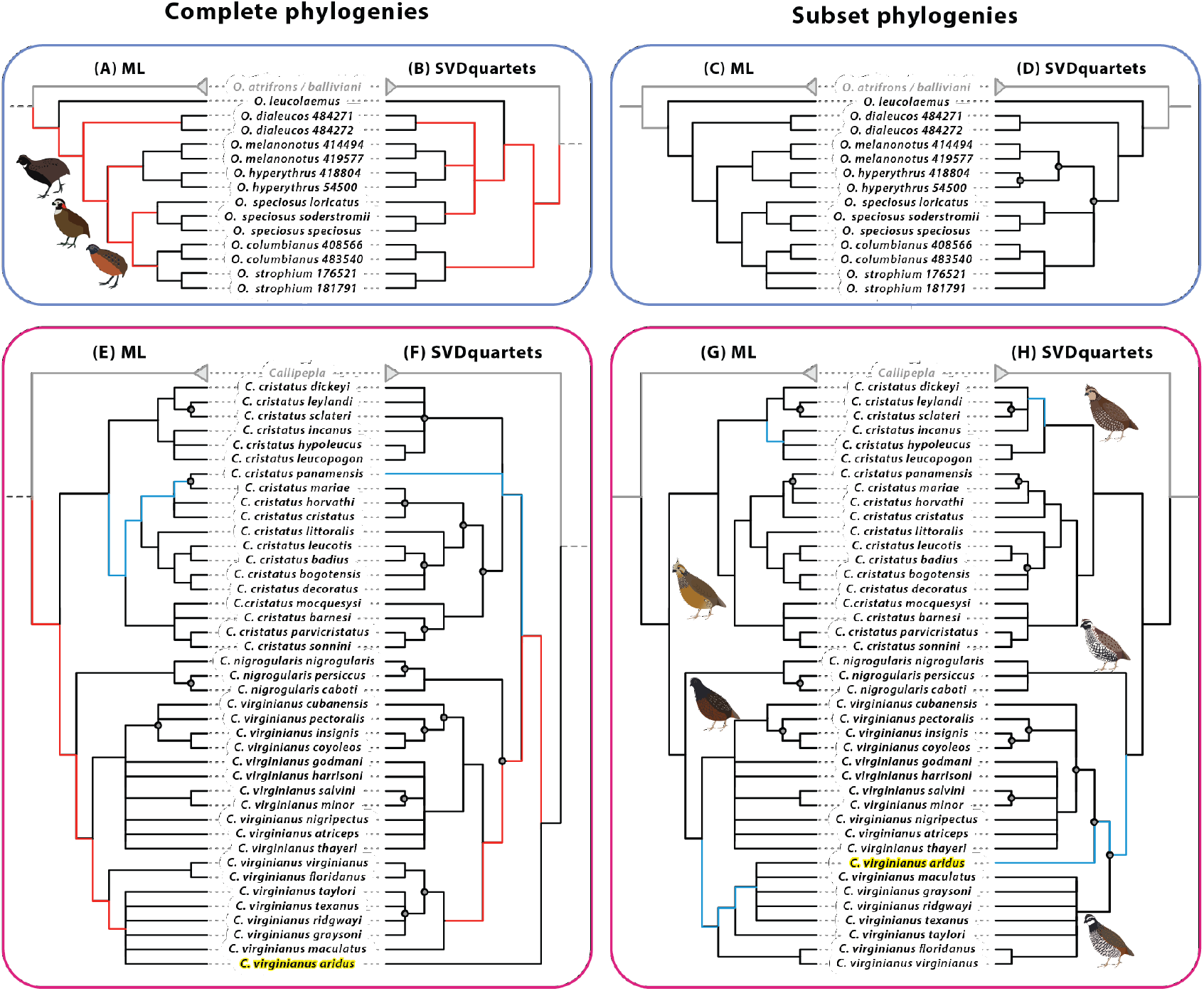
Cladogram of discordant subspecies-level relationships inferred with maximum likelihood (ML) analysis and SVDquartets analysis of 3,884 nuclear ultraconserved element loci within **(A-D)** *Odontophorus* and **(E-H)** *Colinus*. Left panels **(A, B, E, F)** zoom in on the relationships inferred for these clades in analyses of the complete 131 taxon dataset (Fig. 3, Supplementary Fig. S3); right panels **(C, D, G, H)** show relationships inferred using only the subset of taxa depicted. Nodes with >95% bootstrap support are unlabeled; gray circles = 70-94% bootstrap support; nodes with <70% bootstrap support have been collapsed to polytomies. Red branches indicate conflicting species relationships; blue branches indicate conflicting subspecies relationships. Note position change of *C. virginianus aridus* (highlighted in yellow) between complete and subset phylogenies. For branch lengths and the uncollapsed ML tree of all taxa, see Supplementary Fig. S1; for the uncollapsed SVDquartets tree of all taxa, see Supplementary Fig. S2.

### Coalescent UCE phylogeny

The tree we inferred with SVDquartets was well resolved, strongly supported, and largely congruent with the topology we reconstructed with the ML analysis, particularly at the species level (Fig. 3B; see Supplementary Fig. S2 for the uncollapsed topology). In particular, the SVDquartets tree improved support for the sister relationship between the two major *Odontophorus* clades (lowland tropical forest + montane-associated species). At the species level, only two areas of the coalescent tree disagreed with the ML topology: among the seven species in the montane-associated *Odontophorus* clade and within *Colinus*. Within the montane *Odontophorus* clade, the SVDquartets analysis resolved *O. strophium + O. columbianus* as sister to remaining lineages in the clade and suggested a sister relationship between *O. dialeucos* and *O. speciosus*, although this relationship was poorly supported. In contrast, the ML analysis resolved the Central American species *O. leucolaemus* and *O. dialeucos* as successive sister lineages to the remaining sister clades of trans-Andean species (*O. melanononotus + O. hyperythrus*) and cis-Andean species (*O. speciosus, O. strophium* + *O. columbianus*). Within *Colinus*, the major difference was that the SVDquartets analysis resolved *C. virginianus aridus* of northwestern Mexico as sister to all other taxa within *Colinus*.

We observed more discordance between the ML and SVDquartets topologies at the subspecies-level, although most of this discordance existed in parts of either tree having low support (Supplementary Fig. S3). In general, the SVDquartets analysis was less precise, and the lower support values collapsed to a number of polytomies, as seen in *Cyrtonyx*, *Callipepla californica*, and *Dendrortyx macroura* (Supplementary Fig. S3). Despite these areas of low support, the ML and SVDquartets topologies largely agreed in the arrangement of subspecies groups within highly polytypic species, such as *C. cristatus* and *C. virginianus* (Fig. 4).

### Subset concatenated and coalescent phylogenies

For the two clades in which we observed species-level discordance between the ML and SVDquartets trees inferred from the complete dataset, these conflicts were largely resolved when we inferred trees using only those subsets of taxa (Fig. 4). In the *Odontophorus* group, the ML and SVDquartets trees inferred using subclade data both resolved a branching order consistent with the ML topology inferred using the complete dataset, placing the Central American species *O. leucolaemus* and *O. dialeucos* as successive sister lineages to the clade comprising three South American groups, including *O. strophium + O. columbianus* (Fig. 4C-D). Although the subset SVDquartets tree could not resolve the polytomy between *O. hyperythrus + O. melanonotus, O. speciosus,* and *O. strophium + O. columbianus*, this topology was consistent with both ML trees and resolved the major discordance observed in the complete SVDquartets tree, which initially suggested *O. strophium + O. columbianus* were sister to other taxa within this group (Fig. 3B, Fig. 4B).

Within the *Colinus* clade, there were two major differences between the ML and SVDquartets trees inferred with the complete dataset: (1) the placement of *C. cristatus panamensis* in a polytomy in the SVDquartets tree; and (2) the placement of *C. virginianus aridus* sister to all remaining *Colinus* in the SVDquartets trees (Fig. 3B, Fig. 4F). Both the ML and SVDquartets trees inferred with subclade data resolved the *C. [cristatus] leucopogon* clade as sister to the two South American clades of *C. cristatus*, and placed *C. c. panamensis* within one of these South American clades, consistent with the complete ML tree (Fig. 4G-H). Within *C. virginianus*, the ML subset tree inferred subspecies relationships consistent with the complete ML tree, including sister clades of subspecies from southern Mexico and subspecies from northern Mexico + USA, and the placement of *C. v. aridus* within the northern clade of subspecies (Fig. 4G). The subset SVDquartets tree also resolved northern and southern clades of subspecies, consistent with both ML trees, although it placed *C. v. aridus* as sister to the clade of subspecies from southern Mexico (Fig. 3H). Although this placement differs from the ML topologies, the subset analysis resolved the implausible placement of *C. v. aridus* in the complete SVDquartets tree and the major discordance between the SVDquartets and ML trees inferred with the complete dataset (Fig. 4, Fig. 3E-F).

## Discussion

### High-level odontophorid phylogeny is stable across studies, methods, and data types

Across all analyses, we resolved topologies that are consistent with previous studies of major odontophorid clades (Cohen et al., 2012; Crowe et al., 2006; Hosner et al., 2015). Our results from thousands of nuclear loci support the sister relationship between the African genus *Ptilopachus* and the remaining New World odontophorids (Cohen et al., 2012; Crowe et al., 2006; Hosner et al., 2015). Similarly, within the New World clade, we resolved the monotypic genus *Rhynchortyx* as sister to the *Odontophorus* group and the *Dendrortyx* group, a pattern that is generally consistent with previous phylogenetic hypotheses based on morphology and ecology (Johnsgard, 1973). We included representatives of all currently recognized odontophorid species and most analyses inferred consistent, highly-supported relationships at the species level with two notable differences between our UCE topologies, which we discuss below.

Due to a lack of contemporary genetic material, previous molecular phylogenies of odontophorids included just eight of the fifteen described species in *Odontophorus*, the most species-rich genus in the family, with different analyses inferring different topologies for five of the sampled species (Hosner et al., 2015). By including historical specimens, we were able to sample 36 individuals representing all 15 species and all 20 described subspecies (29 taxa in total) for complete taxonomic sampling of *Odontophorus* (Dickinson and Remsen, 2013), and our results across all analyses suggest there are three main lineages within the genus: (1) the northernmost species, *O. guttatus*; (2) a clade of five predominantly lowland tropical forest species; and (3) a clade of nine montane-associated species. These results are consistent with a coalescent topology inferred from a combined mito-nuclear dataset for eight species (Hosner et al., 2015), and with the biogeographic hypothesis that *Odontophorus* had a Central American ancestor (ca. 5.8 Ma) that colonized South America multiple times and diversified rapidly following closure of the Isthmus of Panama (Hosner et al., 2015).

Because we sampled all species, our results refute previous hypotheses that grouped *Odontophorus* species by plumage (Johnsgard, 1988), and they highlight the recurrence of different plumage elements in multiple, presumably independent, radiations – suggesting a shared genomic framework underlying the “mix-and-match” appearance of the 29 taxa in this genus. Johnsgard (1988) recognized three species groups within *Odontophorus* based on shared plumage themes: dark-backed species with rufous fronts (*O. hyperythrus, O. melanonotus*, *O. speciosus,* and *O. erythrops/melanotis*); species with prominent crests and chestnut plumage lacking a white throat (*O. capueira, O. stellatus, O. gujanensis*, and *O. balliviani*); and dark-backed species with striking black-and-white throats and facial patterns (*O. atrifrons, O. leucolaemus, O. dialeucos, O. strophium,* and *O. columbianus*). Our results support some of these relationships, but our results also highlight that similar plumage patterns exist between non-sister lowland and montane species, as well as identifying several previously overlooked plumage similarities between species that our results suggest are closely-related. For example, we resolved a sister relationship between two highly disjunct species, *O. atrifrons* and *O. balliviani*. *Odontophorus atrifrons* is extremely range restricted, inhabiting the subtropical montane forests of northern Colombia and northwestern Venezuela, whereas *O. balliviani* is more broadly distributed throughout montane subtropical forests in southeastern Peru and northern Bolivia (Johnsgard, 1988). Although Johnsgard characterized these species as belonging to two distinct plumage groups, they share a rufous crown and distinctive white, diamond-shaped streaking across the chest. Similar disjunct distributions are observed within other Andean bird species, such as Golden Grosbeaks (*Pheucticus chrysogaster*) (Brewer, 2020) and Red-rumped Bush-Tyrants (*Cnemarchus erythropygius*) (Schulenberg and Kirwan, 2020), and may reflect a history of extinction in the intermediate populations.

### Discordant UCE topologies are artifacts of low-quality historical samples

Our dataset is composed of 66% historical samples (Table 1), and we observed many of the sample quality issues noted in previous studies incorporating this type of material (Moyle et al., 2016; Oliveros et al., 2019; Salter et al., 2020; Swanson et al., 2019). Fortunately, we only recovered species-level topological conflicts in two clades: *Odontophorus* and *Colinus*.

Within both *Odontophorus* and *Colinus*, we observed a previously-noted pattern of discordance that we refer to as the “toepad effect”: in SVDquartets analyses, low-quality samples often aggregate as sister to all other members of the clade in which concatenated analyses place them (Moyle et al., 2016; Oliveros et al., 2019; Salter et al., 2020). By all metrics, the four taxa (represented by six historical samples) that showed this pattern in our analyses (*O. strophium, O. columbianus, C. virginianus aridus,* and *C. cristatus panamensis*) were among the lowest quality historical samples in our dataset: all six samples fell below the 95% confidence interval for cleaned read pairs, number of UCEs, and average contig length (Table 1, Table 2). Three of these samples were collected during the early 1920s, placing them among the oldest samples we sequenced (median collection year 1954), and although the remaining three samples were collected between 1947-1952, specimen preparation and long-term storage conditions can have significant impacts on DNA degradation and quality in addition to sample age (Hall et al., 1997; McCormack et al., 2017; Wandeler et al., 2007).

We further examined the causes of this discordance by analyzing subsets of these data to assess whether spurious relationships could be resolved by reducing the noise to signal ratio introduced with the inclusion of distantly-related taxa. The results of our subset-based phylogenies provide compelling evidence that in our dataset, these “toepad effects’’ are artifacts of sample quality, rather than biological signal. In *Odontophorus*, the concatenated and coalescent subclade trees resolved the discordance we observed among the topologies inferred with the complete dataset, and the subclade topologies ultimately supported the relationships we observed in the complete ML tree (Fig. 4A-D). Although we observed differences in bootstrap support for relationships within *Odontophorus*, such as the polytomy between *O. hyperythrus + O. melanonotus, O. speciosus,* and *O. strophium + O. columbianus* in the subset SVDquartets tree, these differences do not change the relationships or their phylogeographic interpretation within this group (Fig. 4C- D).

Within *Colinus*, the subset topologies resolved the polytomy of *C. cristatus panamensis*, the Central American *C. [cristatus] leucopogon* clade, and the South American *C. cristuatus* clade that we observed in the complete SVDquartets tree, and inferred a placement of *C. cristatus panamensis* consistent with the complete ML tree (Fig. 4E-H). Although the discordant placement of most low-quality samples in our dataset was ameliorated by the subset analyses, some combination of missing loci, exceptionally short contigs, and perhaps other DNA damage proved particularly recalcitrant for *C. virginianus aridus*. Though much improved from the complete SVDquartets tree, the placement of *C. v. aridus* in the subclade SVDquartets tree differs from both ML trees (Fig. 4E-H). Based on the original description of *C. v. aridus* as an intermediate form between *C. v. texanus* and *C. v. maculatus* and its distribution between these two subspecies (Aldrich, 1942), the topology inferred in the ML trees is consistent with our expectations of the relationships among these subspecies; however, we were unable to completely resolve the placement of this sample due to poor data quality.

Our results also underscore the importance of sampling multiple historical specimens within each taxon, when possible. For example, because we resolve the sister relationship of *O. strophium + O. columbianus* across all analyses, we are confident that this relationship is not a “toepad effect”, but likely reflects biological signal and confirms previous hypotheses of a close relationship between these species based on plumage (Johnsgard, 1988). Whereas previous examples of the “toepad effect” have been noted with a single sample per taxon, our results suggest that including multiple toepad samples per taxon can help distinguish between the effects of low-quality samples (as in *C. v. aridus*) and true phylogenetic signal. Considering our results, we advocate for an integrative approach to examining the causes of topological discordance as large datasets encompassing samples of heterogeneous quality become commonplace in phylogenomics.

### Odontophorid taxonomy is largely congruent with genetic data

The impressive phenotypic diversity among odontophorids, especially in male plumage, has contributed to historical fluctuations in odontophorid taxonomy, especially at the subspecies-level (Fig. 2). However, both our ML and coalescent phylogenies using UCEs demonstrate that current taxonomy is largely consistent with the genetic relationships within and among most species of odontophorids (Fig. 3), although we did find two examples of species that were not monophyletic. All analyses (Supplementary Fig. S3) failed to recover *Cyrtonyx montezumae* and *C. ocellatus* as reciprocally monophyletic, instead suggesting these taxa form a grade from north to south. In our concatenated UCE ML tree, the three northernmost *C. montezumae* subspecies form one clade, sister to a clade of *C. ocellatus* and the two Oaxacan subspecies, *C. m. rowleyi* and *C. m. sallei* (Supplementary Fig. S1). Although the SVDquartets analysis recovered a different topology (Supplementary Fig. S2), it still did not support the reciprocal monophyly of *C. montezumae* and *C. ocellatus*, suggesting that population-level sampling and further investigation of plumage, morphology, and vocal data are needed to assess species boundaries within this genus. Based on the available evidence, our results support merging *C. montezumae* and *C. ocellatus* into *C. montezumae* (Vigors, 1830). Similarly, neither our ML or SVDquartets analyses resolved *Odontophorus melanotis* and *O. erythrops* as reciprocally monophyletic (Supplementary Material S3), suggesting these taxa constitute a single species (*O. erythrops,* Gould, 1859), consistent with previous classifications (Johnsgard, 1988). Both of these examples highlight the importance of complete taxon sampling for accurate systematic analysis of polytypic species.

In birds, subspecies designations have traditionally been used to describe diagnosable geographic differences among populations in some morphological or behavioral trait, such as plumage color or song (Mayr, 1982). These subspecies may come into contact in some part(s) of their range, and there is the presumption that gene flow would occur wherever populations come into contact. If gene flow is occurring, this begs the question of how well subspecies relationships can be resolved in a strictly bifurcating phylogenetic framework (reviewed in Phillimore and Owens, 2006), especially with highly conserved genetic markers such as UCEs (Harvey et al., 2016; Smith et al., 2014). Although we could not resolve all subspecies relationships, we were pleasantly surprised by the concordance of well-resolved geographic structure within most polytypic species across all analyses. For example, although the relationships among the eight *O. gujanensis* subspecies differed slightly between the ML and SVDquartets trees (Supplementary Fig. S3), all analyses resolved three groups consistent with the major geologic provinces of the region (Silva et al., 2019): a Central American / northeastern Colombian group (*O. g. castigatus* and *O. g. marmoratus*); a group from west of the Negro and Madeira rivers in the Amazonian foreland basins (*O. g. buckleyi*, *O. g. medius, O. g. rufogularis*, and *O. g. pachyrhynchus*); and a group from east of the Negro and Madeira rivers in the Guiana and Brazilian shields (*O. g. gujanensis* and *O. g. simonsi*) (Supplementary Fig. S3). We observed a similar pattern in *Dactylortyx thoracicus*, for which we sampled nine of the eleven mostly allopatric subspecies, and our analyses consistently resolved the three taxa found north of the Isthmus of Tehuantepec (*D. t. devius, D. t. melodus,* and *D. t. pettingilli*) as sister to a group comprising three pairs of geographically adjacent sister taxa found south of the Isthmus: *D. t. chiapensis + D. t. dolichonyx* from Chiapas; *D. t. sharpei + D. t. paynteri* from the Yucatan peninsula; and *D. t. fuscus + D. t. conoveri* from Honduras (Supplementary Fig. S3). In contrast to these patterns, our results also highlighted several polytypic species with little discernible structure across analyses, such as among the five subspecies of *Oreortyx pictus* or *Callipepla california*, suggesting a review of subspecies designations in these species is warranted.

The power of thousands of genome-wide loci to resolve geographic structure among shallow evolutionary lineages is exemplified in our results for *Colinus*. The three species of bobwhites in the genus *Colinus* epitomize many of the extremes and challenges of odontophorid diversity and taxonomy: together, these three species comprise 44 subspecies described by differences in male plumage, half of which belong to *C. virginianus* (Dickinson and Remsen, 2013). The evolutionary relationships among subspecies within *Colinus* remain largely unclear (Ellsworth et al., 1989; Eo et al., 2009; Evans et al., 2009; Williford et al., 2016, 2014), potentially due to the recent origin of this genus (∼5 MA; Hosner et al., 2015; Williford et al., 2016) and the limited power of the few genetic markers surveyed in prior studies. In contrast to these previous studies, we found strong evidence of geographic population structure within all three species.

Across all analyses, our results suggest the nineteen subspecies of *C. cristatus* compose three well-supported clades, consistent with previous analyses of mitochondrial data (Williford et al., 2016): (1) the six-subspecies *leucopogon* group from Central America; (2) the nine-subspecies *cristatus* group ranging from eastern Panama to the west slope of the Eastern Cordillera and the Caribbean slope of Colombia and Venezuela; and (3) the four-subspecies *sonnini* group ranging from the east slope of the Eastern Cordillera to the Guiana Shield. In contrast to a previous mitochondrial analysis, which resolved the *sonnini* group as sister to *leucopogon + cristatus* (Williford et al., 2016), both our complete and subclade ML and SVDquartet analyses suggest the *leucopogon* group is sister to the remaining subspecies that comprise the *sonnini* + *ciristatus* group (Fig. 4E-H). Species limits within *Colinus* have long been debated (see Johnsgard, 1988), and although these data are insufficient to make taxonomic recommendations for this complex group, we note that dating analyses estimate the deepest divergences within *C. cristatus* are consistent with the timing of divergence between *C. nigrogularis* and *C. virginianus* (∼2.5 MYA; Williford et al., 2016), suggesting species limits have been applied inconsistently across the genus.

Our results also shed new light on the contentious relationships within *C. virginianus*, the most polytypic odontophorid. Historically, 24 subspecies of *C. virginianus* have been described by male plumage (Carroll, 2019), of which 22 are currently recognized (Dickinson and Remsen, 2013), and we collected genomic sequence data from 19 of them. We did not include samples of *C. v. marilandicus* and *C. v. mexicanus*, because these subspecies are often synonymized within *C. v. virginianus*, and we did not include samples of *C. v. nelsoni*, which is often synonymized within *C. v. insignis* (Carroll, 2019). Due to its significance as a game bird in the U.S. and Mexico, *C. virginianus* is one of the most intensively studied bird species (Guthery, 1997), yet previous efforts to understand the relationships among subspecies have yielded equivocal results (Ellsworth et al., 1989; Eo et al., 2009; Evans et al., 2009; Williford et al., 2016, 2014), often finding little evidence of genetic structure. Two possible explanations for findings of panmixia within *C. virginianus* are its recent Pleistocene origin (∼1.5 MYA; Hosner et al., 2015; Williford et al., 2016) and the long history of human-mediated translocations within this species (Whitt et al., 2017). Previous studies have also relied on few genetic markers, which may be insufficiently powerful to resolve shallow genetic structure (Zarza et al., 2016). This is the first study to use thousands of nuclear loci to assess the relationships within *C. virginianus*, and despite shallow divergences, we recover consistent, well-supported geographic structure across all analyses (Fig. 4E-H, Supplementary Fig. S3).

Both our ML and SVDquartets trees using UCEs recover the deepest divergence within *C. virginianus* to be along Mexico’s Transvolcanic Belt, a known biogeographic barrier for birds and other terrestrial taxa (Marshall and Liebherr, 2000; Morrone, 2010), with eight subspecies forming a northern clade and eleven subspecies forming a southern clade. In contrast to the bold white facial patterning typical of *C. virginianus* males in the northern part of their range, seven subspecies of *C. virginianus* males have nearly to completely black heads and throats, including six subspecies from Oaxaca and Chiapas (*atriceps, coyoleos, harrisoni, insignis, nelsoni,* and *salvini*) and the isolated Sonoran desert subspecies *C. v. ridgwayi*, prompting speculation that all black-throated subspecies are closely related (Aldrich, 1946). However, our results suggest that *C. v. ridgwayi* is most closely related to a group of subspecies from Texas and northern Mexico (Fig. 4E-H, Supplementary Fig. S3), consistent with previous findings of shared mitochondrial haplotypes among these populations (Williford et al., 2016). Although our analyses could not resolve all relationships within the southern clade, they do confirm that black- and white-throated subspecies do not form reciprocally monophyletic clades (Fig. 4E-H, Supplementary Fig. S3), suggesting that black throat color may have been gained or lost multiple times within *C. virginianus*.

Our results across all of *Colinus* demonstrate both the power and limitations of phylogenomics for resolving subspecies relationships. By sampling thousands of genome-wide loci from just a single individual per subspecies, we found strong evidence of geographic structure and differentiation among groups of subspecies where previous studies sampling fewer markers have not, highlighting the need for greater sampling of the genome at a finer population scale to disentangle the complex evolutionary history of this genus and inform possible taxonomic revisions.

## Conclusions

We demonstrate the power of UCE phylogenomics to resolve relationships ranging from family-level to below species-level using comprehensive taxonomic sampling of historical museum specimens. While placements of most historical samples were concordant between concatenated and coalescent analyses, we showed that discordant topologies were artifacts of poor sample quality and could be largely resolved by inferring trees using subsets of only those taxa in discordant clades. Within odontophorids, our results affirm previous findings at the genus-level and provide new resolution of species-level relationships, which are largely concordant with current taxonomy. At the subspecies-level, we demonstrate that UCE phylogenomics can resolve consistent, well-supported geographic structure across analyses in most polytypic species, and we highlight the need for increased population-level sampling in several key species complexes, especially within *Odontophorus* and *Colinus*.

## Acknowledgements

We thank the following institutions and people who provided tissue and toepad loans to make this work possible: American Museum of Natural History (AMNH - Paul Sweet & Tom Trombone); Academy of Natural Sciences of Drexel University (ANSP - Nate Rice); Florida Museum of Natural History (FLMNH - Andy Kratter); Field Museum of Natural History (FMNH - Ben Marks, Shannon Hackett, & John Bates); University of Kansas Biodiversity Research Institute (KU - Mark Robbins); Natural History Museum of Los Angeles County (LACM - Kimball Garrett); Louisiana State University Museum of Natural Science (LSU - Donna Dittmann, Van Remsen, & Fred Sheldon); Moore Laboratory of Zoology (MLZ - James Maley); Museo de Zoología ‘Alfonso L. Herrera’ (UNAM - Adolfo Navarro Sigüenza); National Museum of Natural History (USNM - Christopher Milensky); University of Washington Burke Museum (UWBM - Sharon Birks); Western Foundation of Vertebrate Zoology (WFVZ - René Corado); and the Yale Peabody Museum (YPM - Kristof Zyskowski). Marguerite Poché provided assistance with the lab work.

## Ethics statement

Animal tissues used in this research were collected from museum specimens and are not subject to Institutional Animal Care and Use Committee approval.

## Author contributions

J.F.S., P.A.H., E.L.B., R.T.K. and B.C.F. designed the study. J.F.S, P.A.H., W.L.E.T., J.E.M., E.L.B., and R.T.K. contributed samples. J.F.S. collected and analyzed the data with assistance from B.C.F. J.F.S., R.T.B., and B.C.F. wrote the paper with edits from all authors. B.C.F. and R.T.B. contributed funds and supervised the research.

## Data availability

Raw sequencing reads will be available from the National Center for Biotechnology Information (NCBI) Sequence Read Archive (BioProject PRJNA 777908) upon publication. The PHYLUCE computer code used in this study is available from https://github.com/faircloth-lab/phyluce. Other custom computer code, DNA alignments, analysis inputs, and analysis outputs will be available through Dryad upon publication.

