## Supplementary Figures S1-3 for "Historical specimens and the limits of subspecies phylogenomics in the New World quails (Odontophoridae)"

**This PDF file includes:**

Supplementary Figures S1-3

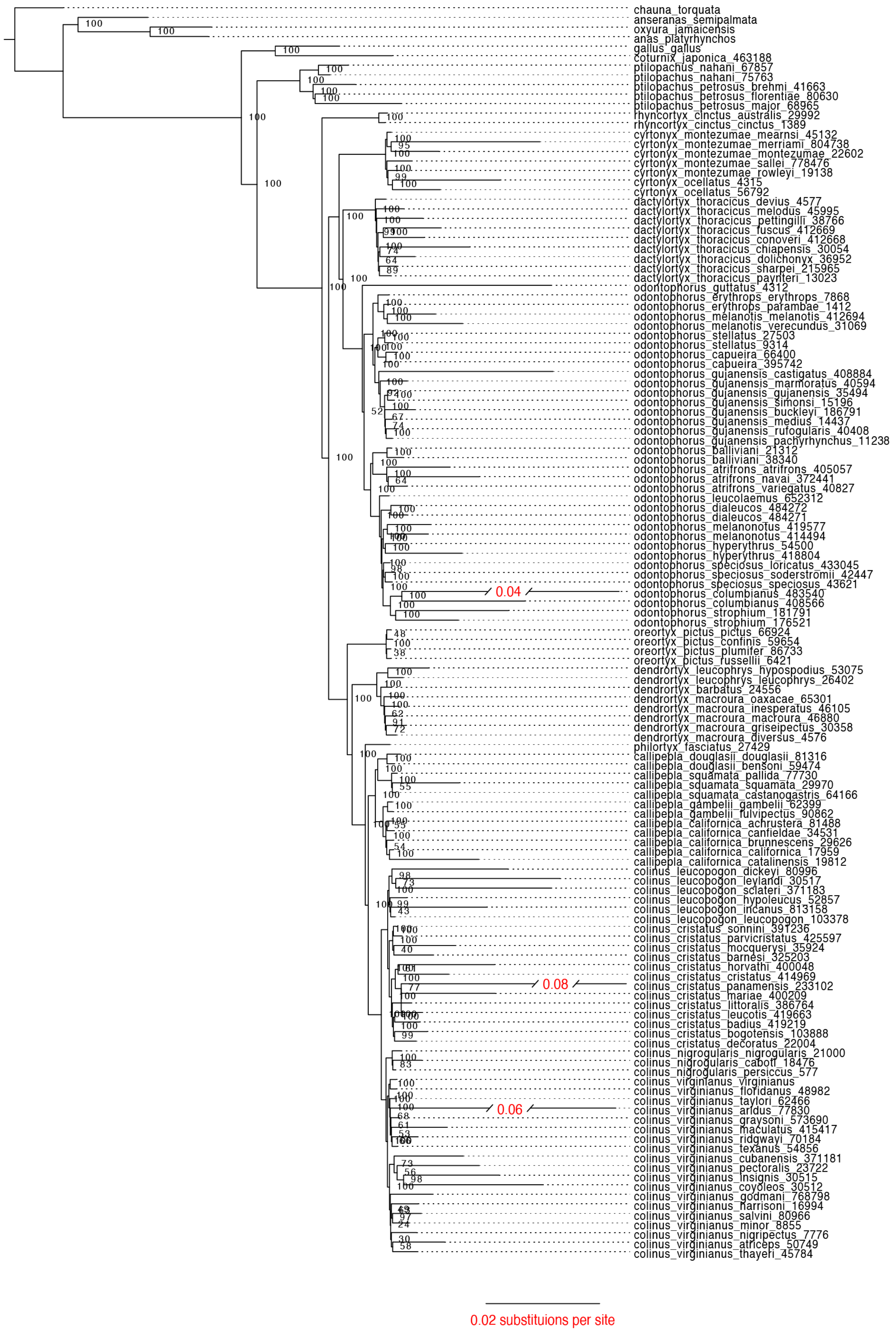


Supplementary Fig. S1: [illegible] 131 taxa (125 [illegible] 6 [illegible]

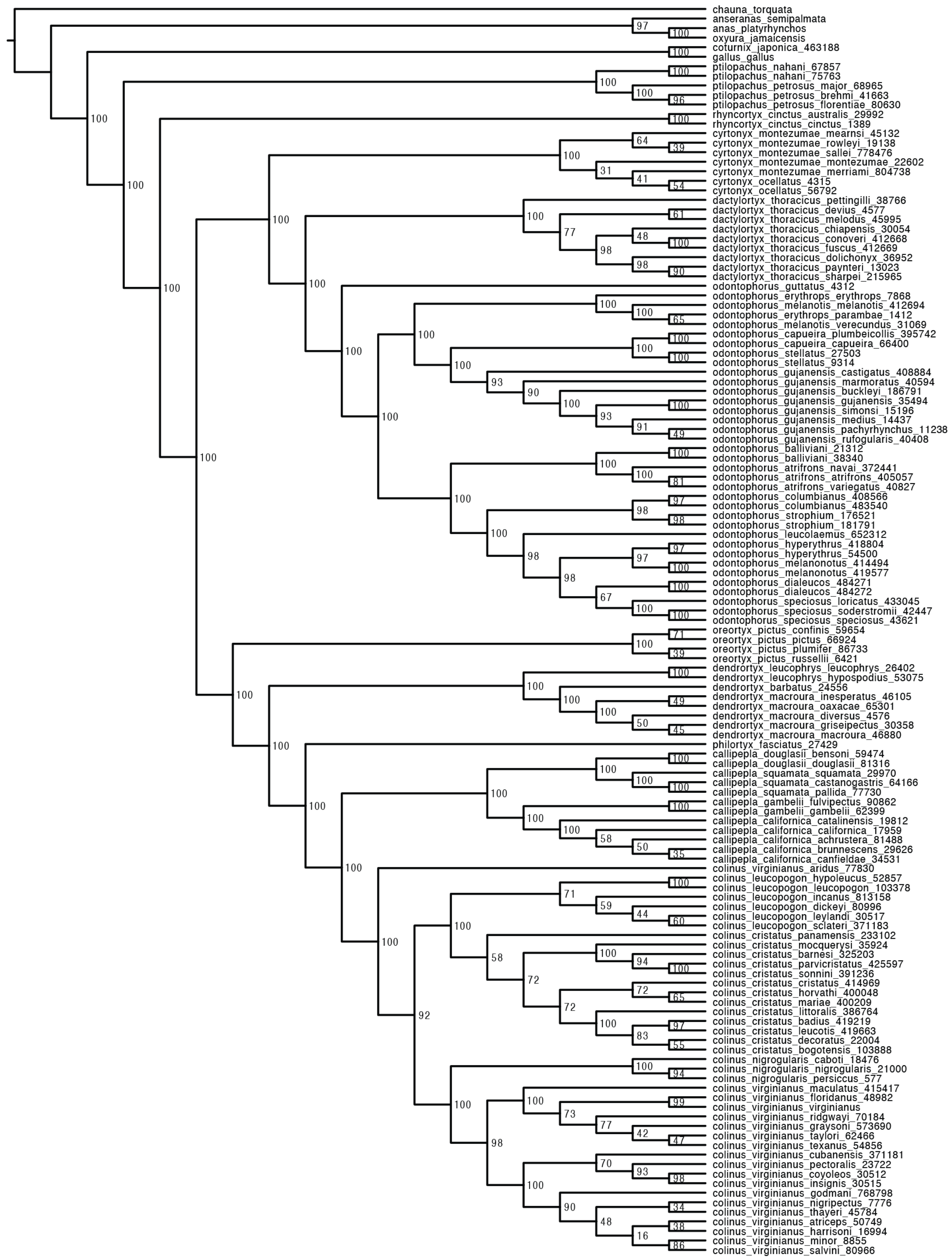


**Supplementary Fig. S2.** Uncollapsed subspecies-level tree of 131 taxa (125 odontophorids and 6 outgroups) inferred with SVDquartets analysis of 3,884 nuclear ultraconserved element loci. Note that SVDquartets does not estimate branch lengths. For a version of this tree in which nodes with bootstrap <70% have been collapsed, see Supplementary Fig. S3.

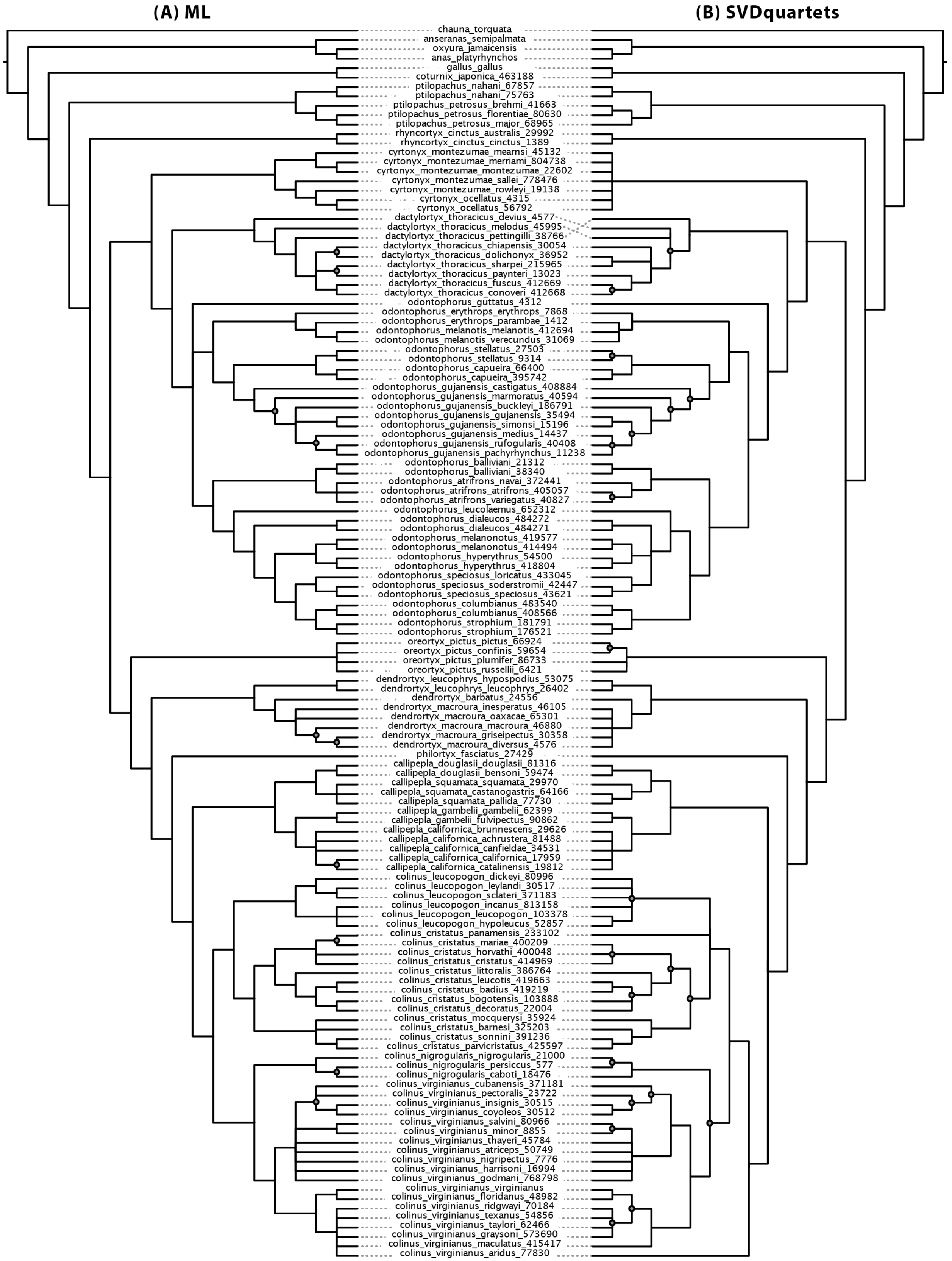


**Supplementary Fig. S3.** Cladogram of subspecies-level relationships of 131 taxa (125 odontophorids and 6 outgroups) inferred with **(A)** maximum likelihood (ML) analysis and **(B)** SVDquartets analysis of 3,884 nuclear ultraconserved element loci. Unlabeled nodes have >95% bootstrap support; gray circles = >70% bootstrap support; branches with <70% bootstrap have been collapsed. For branch lengths and the uncollapsed ML tree, see Supplementary Fig. S1; for the uncollapsed SVDquartets tree, see Supplementary Fig. S2.
